# Particle-based simulations reveal two positive feedback loops allow relocation and stabilization of the polarity site during yeast mating

**DOI:** 10.1101/2023.05.01.538889

**Authors:** Kaiyun Guan, Daniel J. Lew, Timothy C. Elston

## Abstract

Many cells adjust the direction of polarized growth or migration in response to external directional cues. The yeast *Saccharomyces cerevisiae* orient their cell fronts (also called polarity sites) up pheromone gradients in the course of mating. However, the initial polarity site is often not oriented towards the eventual mating partner, and cells relocate the polarity site in an indecisive manner before developing a stable orientation. During this reorientation phase, the polarity site displays erratic assembly-disassembly behavior and moves around the cell cortex. The mechanisms underlying this dynamic behavior remain poorly understood. Particle-based simulations of the core polarity circuit revealed that molecular-level fluctuations are insufficient to overcome the strong positive feedback required for polarization and generate relocating polarity sites. Surprisingly, inclusion of a second pathway that promotes polarity site orientation generated a mobile polarity site with properties similar to those observed experimentally. This pathway forms a second positive feedback loop involving the recruitment of receptors to the cell membrane and couples polarity establishment to gradient sensing. This second positive feedback loop also allows cells to stabilize their polarity site once the site is aligned with the pheromone gradient.

**Author summary:** Cells perform many complex tasks, including directed growth, migration, division and differentiation. To accomplish these tasks, the relevant molecular machinery is localized to specific cellular regions. The asymmetric distribution of cellular components is referred to as cell polarity. Polarity is established by localized activation of the protein Cdc42. Establishing mechanisms that regulate the spatiotemporal activity of Cdc42 is a fundamental area of cell biology. Mating yeast cells dynamically relocate a region of high Cdc42 activity, referred to as the polarity site, and grow toward each other after proper alignment of the sites. We investigated mechanisms that generate dynamic polarity sites by performing particle-based simulations of the biochemical reactions that regulate Cdc42 activity. The reactions contain two positive feedback loops that reinforce Cdc42 activity. The first involves autocatalytic activation of Cdc42 through recruitment of an activator. While necessary for polarity establishment, this feedback loop on its own created a stable polarity site that did not relocate. Incorporation of the second feedback loop, which couples the polarity machinery to extracellular mating signals, generated mobile polarity sites. This feedback loop also provides a mechanism for developing stable alignment of polarity sites. Our findings provide insight into how cells regulate polarity dynamics to accomplish complex tasks.

## Introduction

Cell polarity is the asymmetric distribution of cellular components along some axis. Many cells dynamically adapt the direction of polarization in response to environmental cues. Migratory cells relocate the cell “front” when tracking extracellular signals [1–3]. Pollen tubes, plant root apices, fungal hyphae, neuronal axons, and yeast cells change the direction of polarized growth in response to physical or chemical cues [4–8]. The mechanisms underlying polarity site relocation are not yet understood.

The molecular machinery that controls cell polarity centers on the Rho-family GTPase Cdc42, which is highly conserved among eukaryotes [9,10]. Cdc42 acts as a molecular switch. When bound with GDP, Cdc42 is in its off state and a majority is held in the cytosol through interactions with a GDP dissociation inhibitors (GDIs). Inactive Cdc42 can transition to the membrane and dissociate from the GDI. Membrane-associated Cdc42 is activated by a guanine nucleotide exchange factor (GEF) that promotes the release of GDP to allow binding of GTP. Active GTP-Cdc42 binds various “effector” proteins that regulate the cytoskeleton.

The polarity circuit of the budding yeast *Saccharomyces cerevisiae* has been extensively characterized. Yeast cells polarize their growth during budding and mating. Polarity is established and maintained by autocatalytic positive feedback [11,12]. A cytoplasmic protein complex containing an effector and a GEF associates with active Cdc42 at the membrane and promotes activation of neighboring Cdc42 molecules [11,13]. Because diffusion is slow in the membrane as compared to the cytosol, a cluster of Cdc42 molecules in the membrane is slow to disperse but can recruit cytoplasmic polarity factors from a wide catchment area, sustaining a polarity site. Mathematical models capturing these features recapitulate the patterns of Cdc42 localization observed in vegetative yeast cells: unpolarized cells develop clusters that rapidly coarsen to form a single polarity site that is then stably maintained [12–17]. However, during mating, yeast cells relocate their polarity sites in an apparent search for mating partners [18,19]. This observation raises the question of how the yeast polarity circuit might be modified to allow relocation.

During mating, yeast cells of each mating type express G-protein-coupled receptors (GPCRs) that detect extracellular peptide pheromones released by the opposite mating type [20]. Pheromone binding to the receptor stimulates the Gα subunit to exchange GDP for GTP, which dissociates Gα from Gβγ at the membrane. Gα and Gβγ transmit signals to prepare cells for mating, including activation of pathways that regulate cell polarity. The pathway that allows cell polarity to be influenced by external pheromone gradients is mediated by the scaffold protein Far1, which binds to both Gβγ and the Cdc42 GEF [21–23]. This leads to activation of Cdc42 where there is Gβγ, which reflects the location of active GPCR [22,23]. Thus, an external gradient in pheromone concentration can be translated into an internal gradient in active Cdc42 concentration, enabling a cell to polarize toward a mating partner. However, yeast cells mate in crowded conditions in which global pheromone gradients could be uninformative [24,25], and a shallow gradient may be difficult to interpret given the perturbing effects of molecular noise [26]. Indeed, Cdc42 clusters in mating yeast cells often form at a location that is not directed towards the partner [18,27,28]. Relocation of the polarity site is then needed to successfully mate. During an initial “indecisive phase”, Cdc42 clusters in mating yeast spontaneously appear, disappear, and relocate erratically [18]. This behavior is thought to enable the search for a mating partner, and after the indecisive phase cells develop stable polarity sites oriented towards the chosen partner [29].

What leads to the indecisive behavior of Cdc42 clusters in mating cells? One proposal is that erratic relocation is a product of molecular-level noise acting on the yeast polarity circuit [27,30,31]. Here we confirm that polarity site relocation in a realistic yeast polarity circuit can arise from stochastic noise, but only in a very narrow parameter regime. We find that incorporation of the pheromone-responsive Far1 pathway can dramatically expand the parameter space in which models display polarity site relocation. This same pathway can also explain why polarity eventually becomes stably oriented towards a partner.

## Results

### Noise-driven relocation of the polarity site

To systematically investigate whether the core yeast polarity circuit can yield indecisive Cdc42 clustering like that seen in mating cells, we used a previously validated model developed to describe Cdc42 behavior in vegetative yeast cells [12,17,32,33] (Fig 1A, Table 1). The model species are inactive Cdc42 (which can exchange between membrane and cytosolic compartments), active Cdc42 (which is restricted to the membrane compartment), and a complex between an effector and an activator of Cdc42 (here called Bem1-GEF). This complex also exchanges between membrane and cytosol, and can bind to active Cdc42. At the membrane, the complex promotes activation of inactive Cdc42. The main structural feature of the model is the positive feedback loop formed by mutual activation of Cdc42 and Bem1-GEF (Fig 1A, inset). In this feedback loop, active Cdc42 recruits Bem1-GEF from the cytoplasm and increases its GEF activity. In turn, Bem1-GEF activates Cdc42, leading to further Cdc42 recruitment from the cytosolic pool. This positive feedback loop amplifies local fluctuations in Cdc42 activity eventually leading to the formation of a polarity site [12,34].

**Fig 1.**
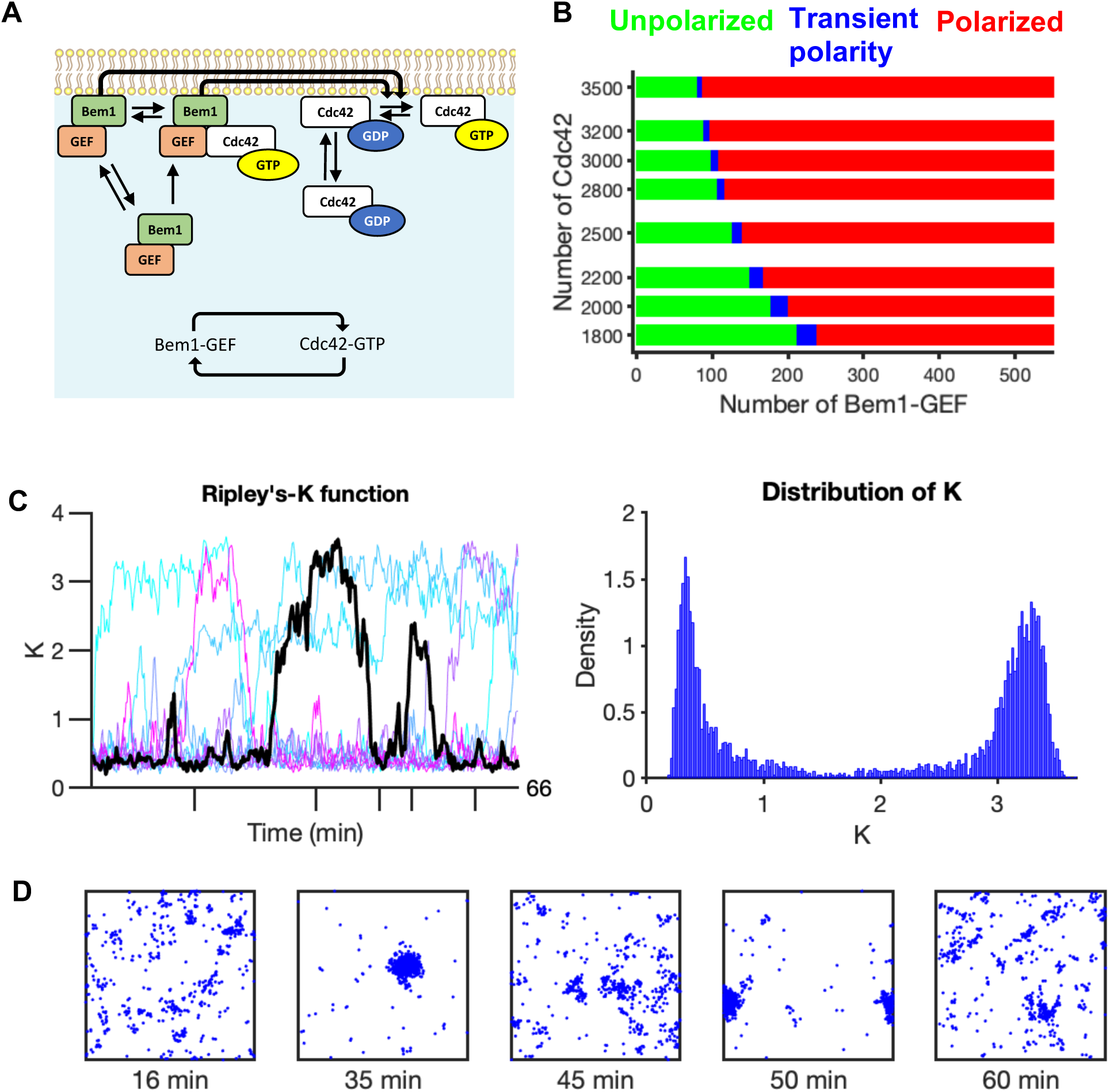
Behavior of the core polarity circuit. **A)** Reaction scheme for the core polarity circuit of yeast (diagram adapted from Pablo et al. [17]). Polarity is driven by positive feedback through mutual activation (Inset). **B)** Two-parameter bifurcation diagram for molecular abundances of Cdc42 and Bem1-GEF (Unpolarized regime – green, transient polarity – blue, polarized – red). **C)** Simulation results for the transient polarity regime. Time series for Ripley’s K-function values (a measure of Cdc42 clustering) were generated for ten simulations (left panel). Distribution of *K* values (right panel). **D)** Distributions of Cdc42-GTP molecules at indicated times. Distributions are taken from the black time series in **C** using time points indicated with ticks on the time axis. Simulations correspond to 3000 Cdc42 and 102 Bem1-GEF molecules using uniform random distributions for Cdc42 and Bem1-GEF as initial conditions.

**Table 1.**
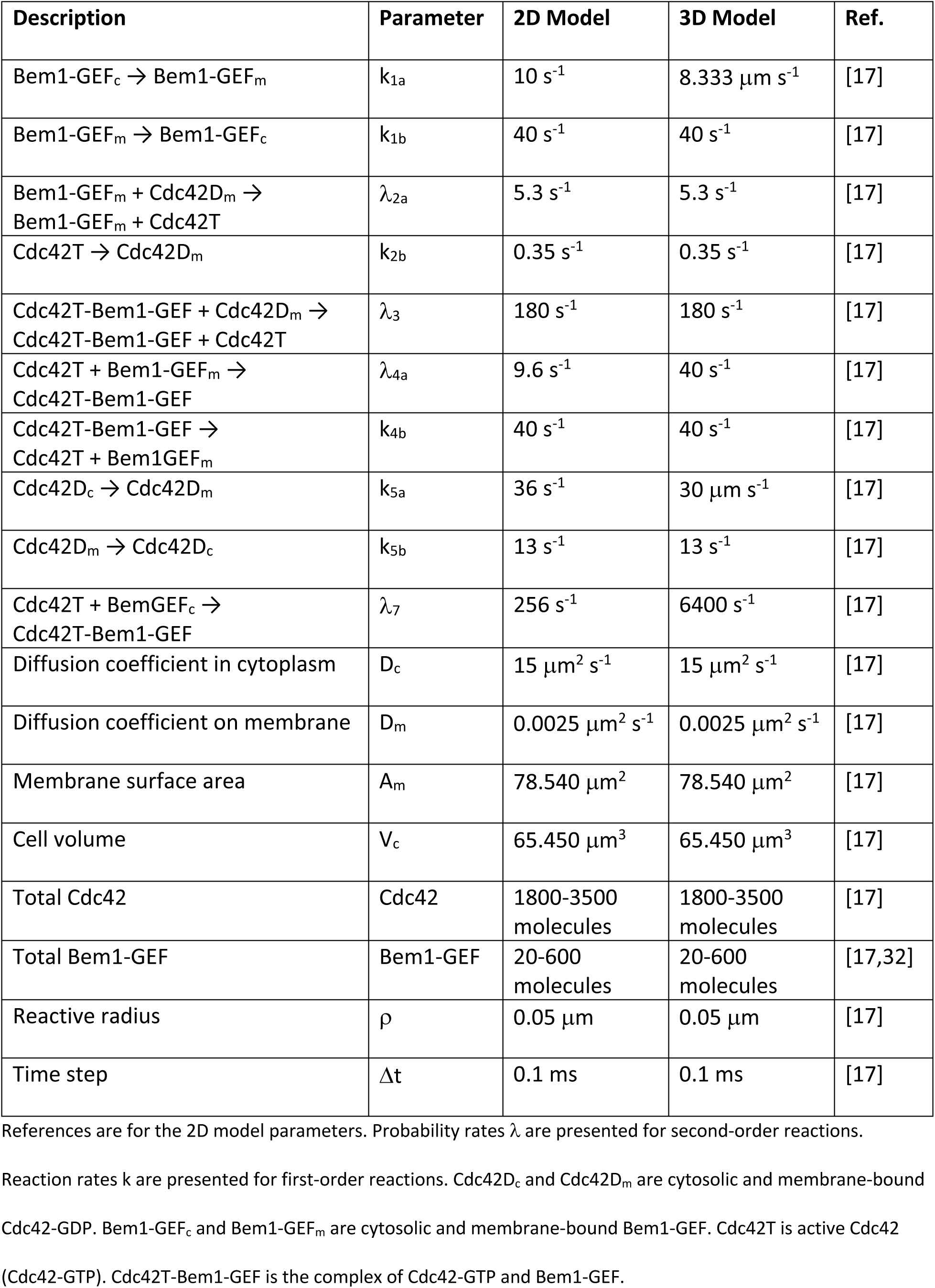
Parameters for the core polarity circuit.

Our first goal was to determine if molecular-level fluctuations arising from the stochastic nature of biochemical reactions and diffusion are sufficient to promote relocation of the polarity site. To test this possibility we used the simulation platform Smoldyn [35,36] to perform particle-based simulations of the core polarity circuit, with parameters described previously [17]. We performed simulations with varying numbers of Cdc42 and Bem1-GEF molecules. These initial simulations revealed three distinct types of behavior: 1) unpolarized, in which no significant Cdc42 clusters formed (S3A Fig), 2) polarized, in which a single strong cluster was maintained (S3B Fig), and 3) transient polarity, in which clusters could spontaneously form, disappear, and re-form (Figs 1C and 1D, S1 Movie). These results suggest that molecular-level noise is able to trigger intermittent transitions between polarized and unpolarized steady states, consistent with previous work on regulatory systems [37–39].

To quantify our results, we used a normalized version of Ripley’s K-function (*K*) [40,41]. *K* measures clustering strength as the deviation of the observed particle distribution from that of a uniform distribution [17]. A value of *K* close to zero signifies a uniform distribution, and the larger the value of *K* the more polarized the distribution. We designated a threshold value of 1.5 to distinguish unpolarized (*K* < 1.5) from polarized (*K* ≥ 1.5) states for all 2D simulations (S1A Fig). In the transient polarity regime, the *K* values exhibit a bimodal distribution, consistent with a bistable system that switches between polarized and unpolarized states (Fig 1C, right panel) [16,37–39]. To investigate how robust the polarity circuit is to transient polarity, we developed a method for distinguishing the three types of behavior based on the distribution of *K* values (Methods and S2 Fig). Our analysis revealed that the parameter regime supporting transient polarity is limited to a small range of molecular abundances (Fig 1B), as also found in earlier work [32]. Thus, it would require very precise control for cells to exploit this parameter regime to develop transitory polarity. Moreover, this model regime exhibits transients between long-lived unpolarized and polarized states, rather than the rapidly fluctuating polarity states observed in indecisive yeast cells. We conclude that the indecisive polarity behavior of mating yeast cells is unlikely to be a simple consequence of molecular noise acting on the core polarity circuit.

### The receptor-Far1 pathway

During mating, there is an additional mechanism for bringing GEF molecules to the membrane. Specifically, following exposure to pheromone, GEF molecules in complex with Far1 (Far1-GEF) are recruited to the membrane through Far1’s interaction with Gβγ released by active receptors. Far1’s interaction with Gβγ couples the polarity circuit to the extracellular pheromone concentration. In addition, the receptor-Far1 circuit forms a second positive feedback loop, because active Cdc42 leads to the trafficking of new receptors to the membrane (Fig 2A). To gain insight into the behavior of the receptor-Far1 circuit, we considered a model consisting of Cdc42, receptors and Far1-GEF, neglecting Bem1-GEF. To simplify the model, the G proteins that mediate interaction between receptors and Far1-GEF are not modeled explicitly. That is, we assume that active receptors recruit Far1-GEF from the cytoplasm. Once at the membrane Far1-GEF promotes Cdc42 activation with the same efficacy as Bem1-GEF in complex with active Cdc42 (Fig 2A, Table 2). We also assume all pheromone receptors are active (this is analogous to the situation in which cells are exposed to saturating pheromone concentration). Pheromone receptors are delivered to the cell surface via Cdc42-oriented actin cables. The model does not explicitly take actin cables or vesicle delivery into account. Rather, we assume the rate at which receptors are delivered is determined by the local concentration of active Cdc42 (see Methods for details). Surface receptors are internalized via endocytosis, at well-characterized rates [42,43]. We further assume that receptors diffuse very slowly at the membrane, consistent with experimental findings [18]. As noted above, because Cdc42 clusters enhance local accumulation of receptors, and receptors recruit Far1-GEF to activate Cdc42, this pathway forms a mutual activation positive feedback loop (Fig 2A, inset). Thus, we anticipated that the receptor-Far1 circuit, like the Bem1-GEF circuit (Fig 1A), would have the capacity to spontaneously polarize.

**Fig 2.**
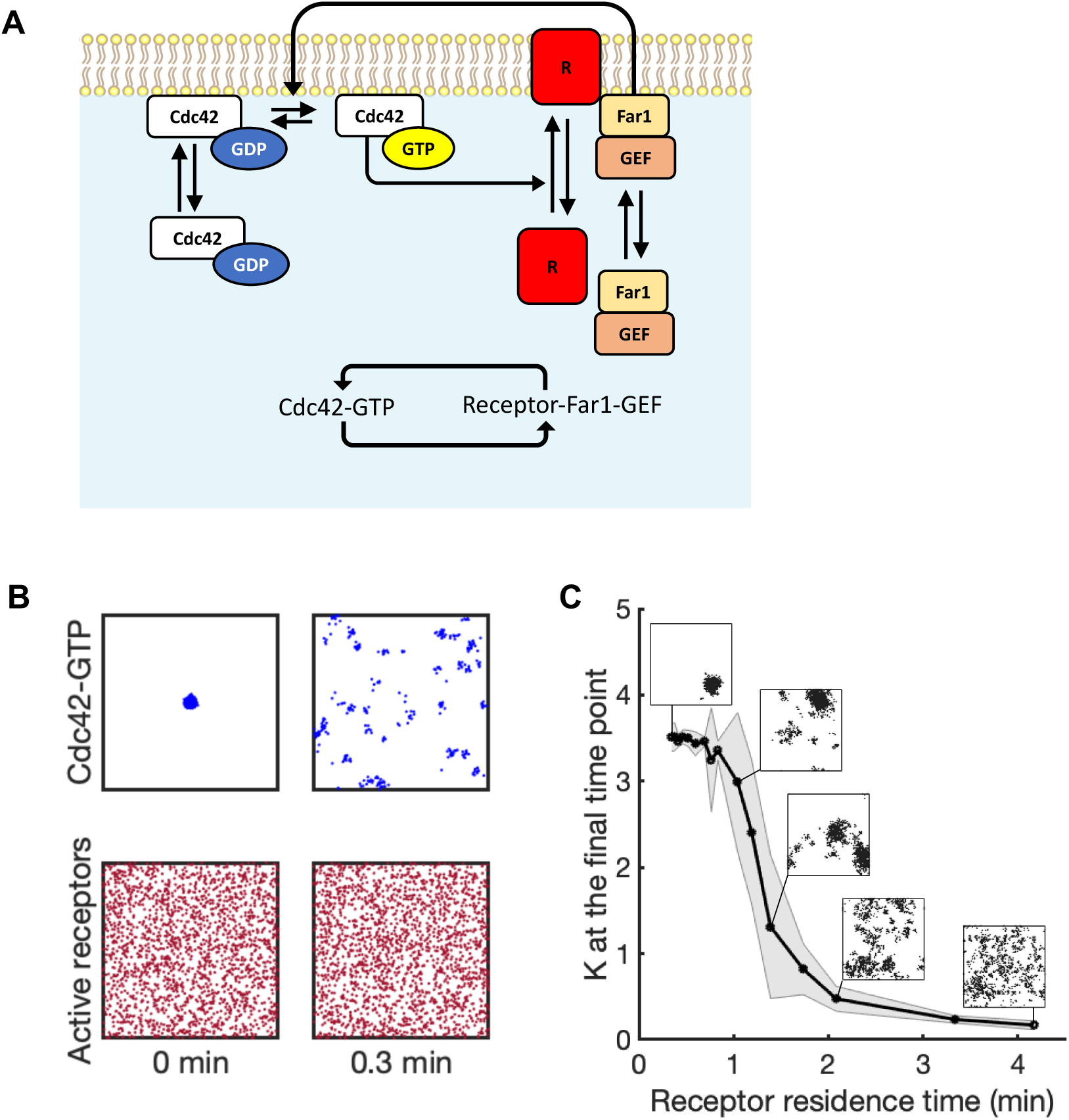
Behavior of the receptor-Far1 circuit. **A)** Reaction scheme for the receptor-Far1 circuit. This circuit forms a second positive feedback loop through mutual activation (inset). **B)** Distributions of active Cdc42 (top row) and receptors (bottom row) shown for 0 min and 0.3 min. Long receptor lifetimes prevent maintenance of a polarity site. Simulations performed using 3000 Cdc42, 280 Far1-GEF, and 2500 receptor molecules. Initial conditions consisted of a polarized cluster of 3000 Cdc42-GTP and uniformly distributed receptor and Far1-GEF molecules. **C)** Final *K* values as a function of receptor membrane residence time. Values of *K* taken at t = 4000 secs. Data points represent averaged values of *K* for 30 simulations and the shaded area denotes standard deviation. Snapshots are for active Cdc42 at t = 4000 secs.

**Table 2.** Parameters for the uniformly activated receptor-Far1 pathway.

| Description | Parameter | 2D Model | 3D Model | Ref. |
| --- | --- | --- | --- | --- |
| $\text{Far1-GEF} + \text{Ra}_m \rightarrow \text{RaGEF}$ | $\lambda_{8a}$ | $300 \text{ s}^{-1}$ | $7500 \text{ s}^{-1}$ | |
| $\text{RaGEF} \rightarrow \text{Far1-GEF} + \text{Ra}_m$ | $k_{8b}$ | $0.11 \text{ s}^{-1}$ | $0.11 \text{ s}^{-1}$ | [44] |
| $\text{Cdc42D}_m + \text{RaGEF} \rightarrow \text{Cdc42T} + \text{RaGEF}$ | $\lambda_{2c}$ | $180 \text{ s}^{-1}$ | $180 \text{ s}^{-1}$ | |
| $\text{Ri}_c + \text{Cdc42T} \rightarrow \text{Ra}_m + \text{Cdc42T}$ | $\lambda_9$ | $0.025 \text{ s}^{-1}$ | $0.625 \text{ s}^{-1}$ | |
| $\text{Ra}_m \rightarrow \text{Ri}_c$<br>$\text{RaGEF} \rightarrow \text{Far1-GEF} + \text{Ri}_c$ | $k_{10}$ | $0.002 \text{ s}^{-1}$ | $0.002 \text{ s}^{-1}$ | [42,43] |
| Diffusion coefficient of RaGEF and $\text{Ra}_m$ | $D_{\text{receptor}}$ | $0.0001 \mu\text{m}^2 \text{ s}^{-1}$ | $0.0001 \mu\text{m}^2 \text{ s}^{-1}$ | [18] |
| Total Far1-GEF | Far1-GEF | 5-280 molecules | 5-280 molecules | [45] |
| Total receptors | Ra + Ri | 2500 molecules | 2500 molecules |  |

We let $\lambda_{2c} = \lambda_3$ to make the GEF activity of Far1-GEF equal to that of Bem1-GEF when bound with Cdc42-GTP. $Ri_c$
and $Ra_m$ are cytosolic and membrane-bound receptors. Receptors are active on the membrane while inactive in the cytosol. RaGEF is the complex of Far1-GEF and $Ra_m$ .

When we simulated this circuit using realistic receptor trafficking rates and initial particle distributions drawn from a uniform distribution, the model did not spontaneously develop a polarized cluster of Cdc42 in biologically relevant timescales. Even when the initial conditions included a strong cluster of active Cdc42 but distributed receptors, the initial clustering was rapidly lost (Fig 2B). To understand why a polarized distribution of Cdc42 was not maintained, consider that receptor trafficking occurs at a much slower rate than the rates for Cdc42 and Far1-GEF association and dissociation: the average membrane residence time of receptors is about 8 minutes [42,43] compared to a few seconds for Cdc42 [46,47] and Far1-GEF [44]. Thus, any clusters of Cdc42 dissipate before they can be reinforced by receptor traffic. Consistent with that interpretation, when the receptor residence time was decreased by increasing receptor trafficking rates the receptor-Far1 circuit did spontaneously establish polarity (Fig 2C).

Our simulations indicate that in spite of its positive feedback loop (Fig 2A), the receptor-Far1 circuit by itself cannot establish a polarity site due to the mismatch between the residence times for Cdc42 (short) and receptor (long). However, it is not clear how the Far1 pathway might influence polarity behavior if the Bem1-Cdc42 circuit was also present, as is the case during the indecisive phase. To understand how these pathways interact, we modeled a system that combined the polarity circuit (Bem1-GEF and Cdc42) with the receptor-Far1 pathway (receptors and Far1-GEF).

### The receptor-Far1 circuit promotes polarity site relocation and generates indecisive behavior

The combined polarity circuit (Fig 3A) includes two interconnected positive feedback loops, one from the core polarity circuit and the other from the receptor-Far1 circuit (Fig 3A, inset). Importantly, these feedback loops operate on different time scales, with the faster polarity circuit supporting polarization, whereas the slower receptor-Far1 circuit is unable to generate a stable polarity site. Therefore, it is not obvious how the combined system will behave. To investigate the behavior of the combined polarity circuit we performed simulations starting from two different initial conditions: a uniform random distribution of pathway components and a pre-polarized Cdc42 cluster. Similar to the polarity circuit alone (Fig 1B), the combined system exhibited one of three behaviors (Fig 3B and S4 Fig) depending on the molecular abundance of the components: an unpolarized regime with little Cdc42 clustering (S5A Fig), a polarized regime with stable and static clusters (S5B Fig), and an intermediate regime where clusters can form or disappear (Fig 3C). However, the unpolarized regime was expanded and the transition to the polarized regime required a larger amount of the Bem1-GEF complex (compare Figs 3B and 1B). In between, there was a substantially larger transient polarity regime. Interestingly, in the transitory regime the behavior of the system appeared to be qualitatively different from the model with only the core polarity circuit. Rather than a single well-formed polarity site appearing and dispersing (Fig 1D), the addition of Far1-GEF seemed to promote multiple transient Cdc42 clusters that dynamically relocate in a manner reminiscent of indecisive polarity clustering in yeast cells (Fig 3C, S2 Movie). To quantify this observation, we employed a cluster detection algorithm based on Voronoi tessellation [48] and tracked the number of detected clusters over time (S6 Fig). In confirmation of our qualitative observations, coexisting Cdc42 clusters in the combined model were more frequent and exhibited more dynamic behavior than in the core polarity circuit alone (compare S6A and S6B Figs).

**Fig 3.**
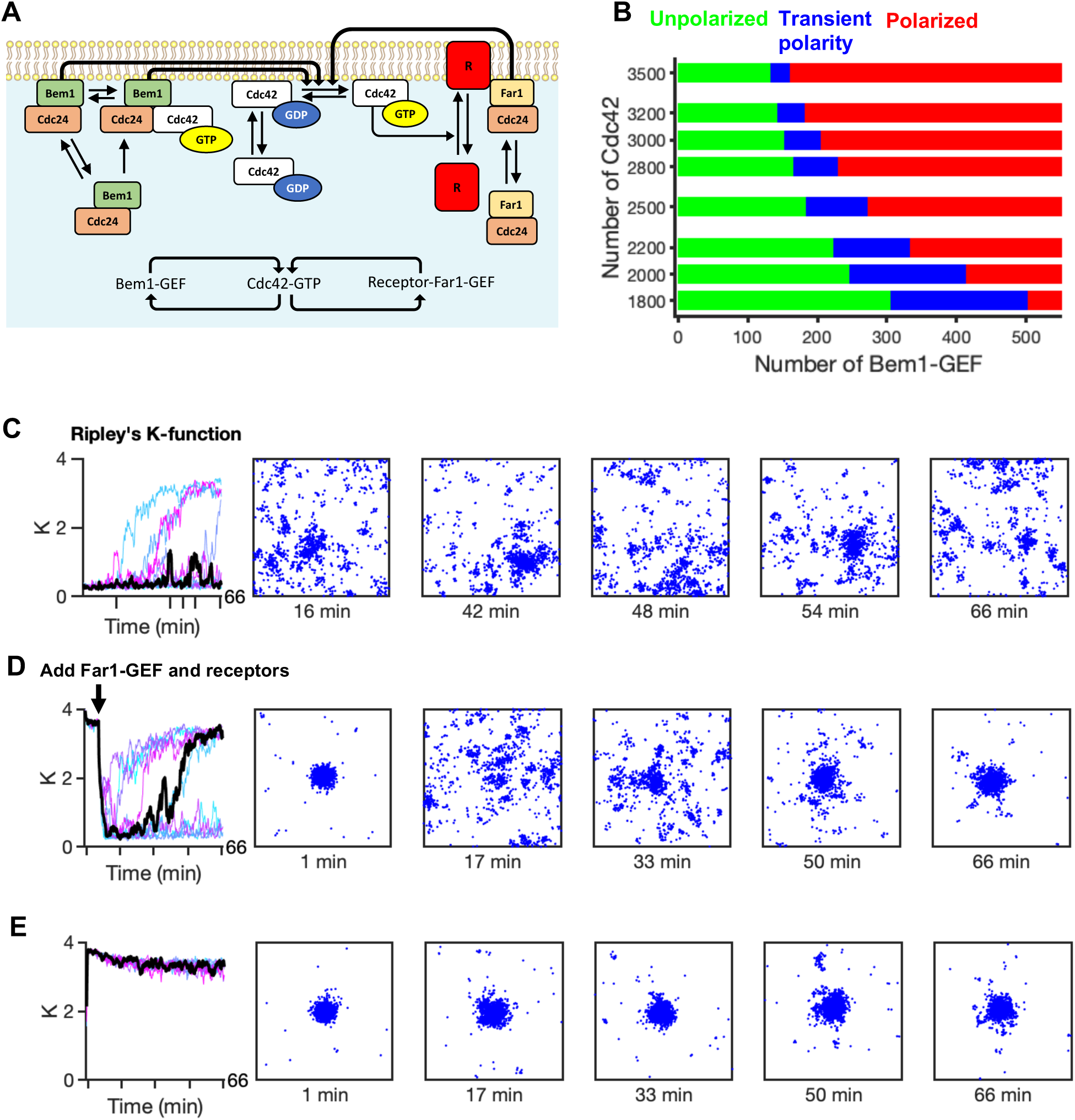
Combined circuit allows relocation and stabilization of the polarity site. **A)** Reaction scheme for the combined polarity circuit. The combined model consists of dual positive feedback architecture (inset). **B)** Two-parameter bifurcation diagram for molecular abundances of Cdc42 and Bem1-GEF (Unpolarized regime – green, transient polarity – blue, polarized – red). **C) – E)**: Simulations results for the transient polarity regime. Plots for *K* consist of 10 realizations. The distributions for active Cdc42 come from the black time series using the time points indicated by tick marks on the time axis. All simulations were performed using 3000 Cdc42, 170 Bem1-GEF, 30 Far1-GEF and 2500 receptor molecules, with different initial conditions as indicated. **C)** Multiple transient polarity sites form using uniform random distributions as initial conditions. In a subset of the simulations, a single polarity site is eventually formed. **D)** An established polarity site rapidly dissipates following the addition of randomly distributed receptors and Far1-GEF molecules. **E)** A polarity site is rapidly established and stably maintained for simulations started with a cluster of active receptors and uniformly distributed Cdc42 molecules.

When we initiated the simulations with a pre-polarized Cdc42 cluster but uniform receptors, the cluster immediately dissipated (Fig 3D, S3 Movie). Thus, even though the receptor-Far1 circuit has a positive feedback architecture, the slow timescale of receptor trafficking allows distributed receptors to overcome Bem1-GEF-maintained polarity. Note that in many of the simulations the polarity site did eventually reform (Fig 3D). In these cases the receptor distribution also became polarized.

We suggest the following mechanism to explain how relocation and indecisive behavior arise in the combined model. Far1-GEF molecules recruited to the membrane by active receptors can seed multiple small Cdc42 clusters, which are then amplified by Bem1-GEF-mediated positive feedback. Without Far1-GEF, clusters would compete to yield a single winning polarity site [16,17]. However, as this competition process begins, Far1-GEF nucleates new clusters, thwarting effective competition: although some Cdc42 cluster(s) may transiently become dominant, newly seeded clusters continue to compete such that no stable cluster can develop. For this mechanism to operate, there are several requirements that the system must satisfy.

First, the distributed nature of active receptors is key to the continual seeding of new clusters: if receptors were instead clustered, then the combined Bem1-GEF and Far1-GEF positive feedbacks would act synergistically to maintain a Cdc42 cluster. This was indeed the case. When simulations were run with an initially focused receptor distribution, all simulations produced a well-polarized Cdc42 distribution (Fig 3E, S4 Movie). As initial receptor distributions were varied, more dispersed receptor distributions lowered the probability that the system would remain polarized (Fig 4A).

**Fig 4.**
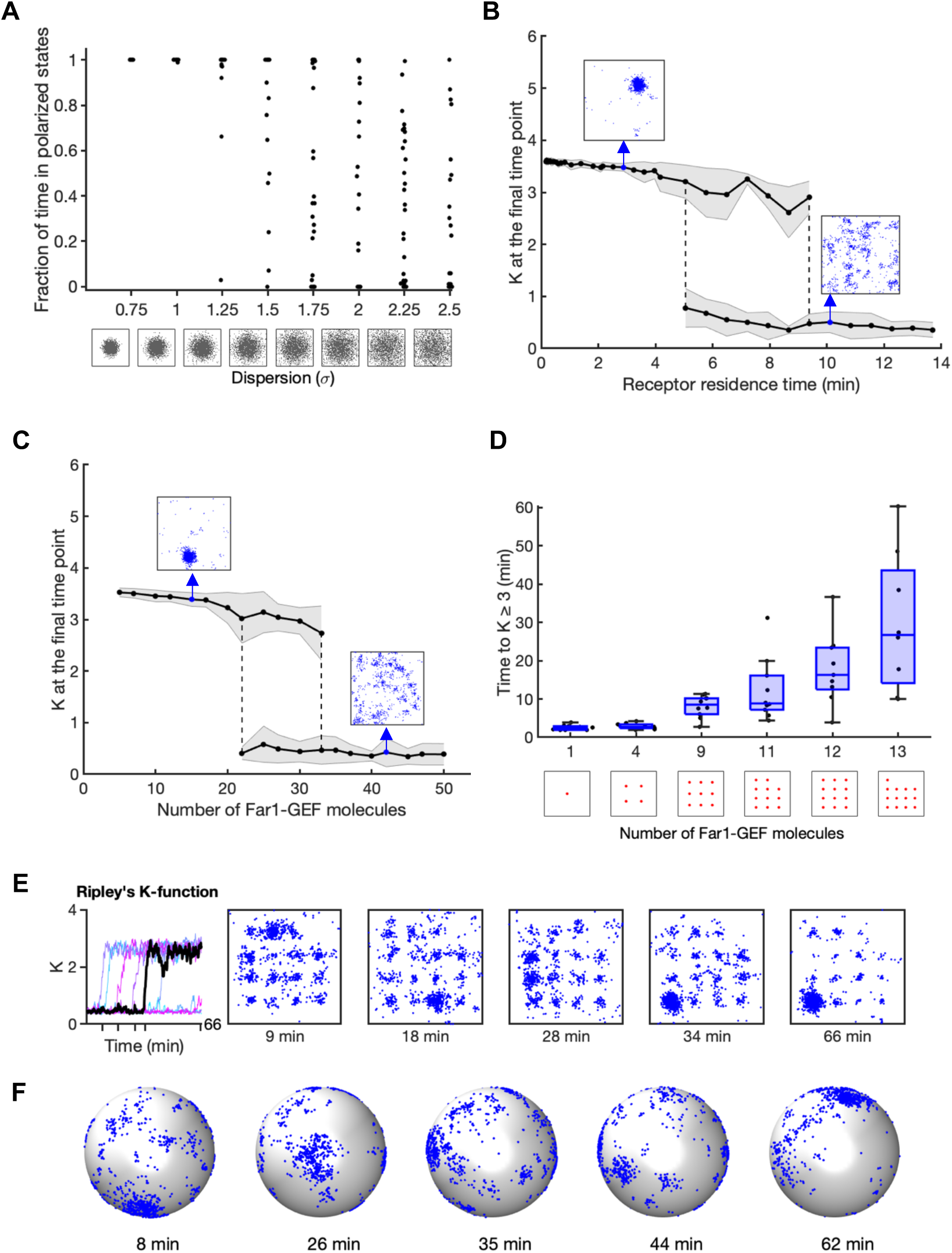
The Far1 pathway promotes indecisive behavior by delaying or preventing coarsening of polarity sites. **A)** The fraction of time polarized as a function of the degree of dispersion in the initial receptor distribution. The fraction of time was measured as the ratio of *K* values ≥ 1.5 to the total number of *K* values in the interval 2000 to 4000 secs to avoid any initial transients. Each point represents a single simulation result from 30 total simulations. Initial receptor distributions were generated from 2D Gaussian distributions. The degree of dispersion represents the standard deviation of the distribution. **B)** Final *K* values (t = 4000 secs) as a function of receptor membrane residence time. 5 ≤ *n* ≤ 30 simulations for each data point. Standard deviations are shaded. **C)** Final *K* values as a function of Far1-GEF molecule number. 5 ≤ *n* ≤ 30 simulations for each data point. **D)** Time to stabilization of the polarity site (K ≥ 3) as a function of number of fixed Far1-GEF molecules in the membrane. Distributions of Far1-GEF molecules are shown along the x-axis. **E)** Time series of *K* values for the case of 15 fixed Far1-GEF molecules (left panel). Right panels show distributions for active Cdc42 molecules for the black time series at times indicated with ticks on the time axis. **F)** Results for a 3D particle-based simulation. Cdc42T molecules shown as blue dots on the cell surface. Rate constants converted from 2D case as described in the Methods and listed in Tables 1 and 2. All simulations performed using 3000 Cdc42, 170 Bem1-GEF, 2500 receptor and unless otherwise noted 30 Far1-GEF molecules.

Second, slow trafficking of pheromone receptors is key to promote indecisive behavior: if trafficking rates were to be increased, then receptor internalization would deprive Far1 of the potential for seeding competing clusters, while delivery of receptors to currently dominant clusters would stabilize them. Indeed, we found that accelerating receptor traffic (reducing receptor residence time) switched the behavior of the combined model, from indecisive relocation to stable polarity (Fig 4B).

Third, increasing the abundance of Far1-GEF would increase the number of newly seeded Cdc42 clusters, driving the system towards a more uniform Cdc42 distribution. This was also observed (Fig 4C). Changing either receptor trafficking rates or Far1-GEF abundance revealed parameter regimes that produced unpolarized states, polarized states, or switching between the two (Figs 4B and 4C).

To directly visualize the hypothesized competition between clusters nucleated by Far1-GEF, we distributed immobile Far1-GEF molecules at specific locations in the simulation domain, so that each fixed Far1-GEF molecule could initiate a cluster of Cdc42. As the number of Far1-GEF molecules increased, the competition time between seeded Cdc42 clusters also increased, delaying the establishment of a single “winner” polarity site (Fig 4D). Cdc42 clusters relocated between different Far1-GEF “seeds” in an indecisive manner, and even when one cluster became dominant, smaller clusters continued to coexist (Fig 4E, S5 Movie).

To confirm that our results held in three dimensions, we performed 3D simulations of the combined polarity circuit. Most parameters for 2D and 3D simulations were kept the same (Table 1-2) so that the steady-state amount of species were similar between the two types of simulations (S7 Fig). Polarity sites in the 3D simulations relocated erratically around the cell cortex, as seen with indecisive polarity clustering in yeast cells (Fig 4F, S6 Movie). In sum, these simulations indicate that a system combining the core Bem1-GEF circuit with the receptor-Far1 circuit can exhibit indecisive polarity behavior, with relocating clusters of Cdc42.

### Pheromone gradients act to stabilize the polarity site

Our simulations demonstrated the Far1 pathway can destabilize polarity to promote relocation in the presence of a uniform pheromone concentration. By contrast, experimental studies found that this pathway is required for orientation and stabilization of polarity sites towards a mating partner [27,49–51]. One way to reconcile these findings is that the Far1 pathway’s effects are dependent on the pheromone concentration profile that cells experience.

A yeast cell emits ∼ 1400 pheromone molecules per second [52]. With this number, the maximum pheromone concentration that the partner cell is exposed to is likely to be below 10 nM [19]. Surprisingly, yeast cells secreting only 20% as much pheromone (peak concentration around 1-2 nM) are still able to stabilize their mating partner’s polarity site [53]. This is well below the concentration of uniform pheromone required for stabilization of a polarity site [51,54]. These findings suggested it may be the pheromone gradient, rather than the absolute pheromone concentration, that stabilizes the position of the polarity site [53]. To test the plausibility of this hypothesis, we included pheromone molecules in our combined polarity circuit simulations (Fig 5A, Table 3). In this situation the receptors are activated upon pheromone binding, allowing us to test whether a pheromone gradient can change polarity site behavior.

**Fig 5.**
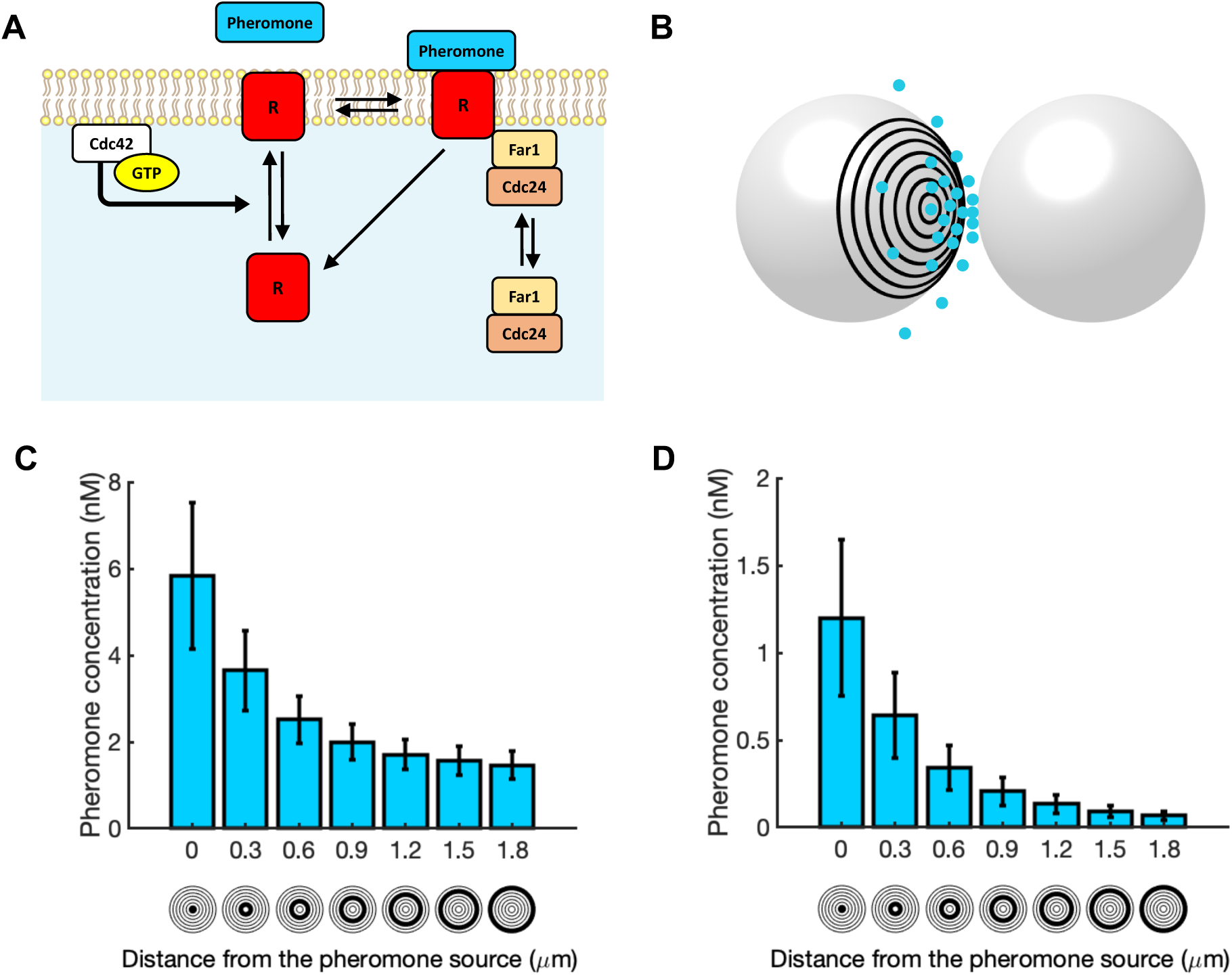
Simulating pheromone gradients. **A)** Reaction scheme for pheromone receptor binding. **B)** A 3D pheromone gradient was simulated between an emitter and receiver cell as described in Clark-Cotton et al. [19]. **C** and **D)** Time-averaged pheromone concentrations for two different emission rates (C - 650 molecules/sec with a uniform background pheromone concentration of 1.5 nM, and D - 150 molecules/sec). Concentrations were measured in the “rings” shown below the x-axis. Error bars denote ± std.

**Table 3.** Parameters for the pheromone-induced receptor-Far1 pathway.

| Description | Parameter | 3D Model | References |
| --- | --- | --- | --- |
| $R_{im} + \text{pheromone} \rightarrow R_{am}$ | $\lambda_{11}$ | $1.8 \text{ s}^{-1}$ | |
| $R_{ic} + \text{Cdc42T} \rightarrow R_{im} + \text{Cdc42T}$ | $\lambda_{12}$ | $0.625 \text{ s}^{-1}$ | |
| $R_{am} \rightarrow R_{im} + \text{pheromone}$ | $k_{13}$ | $0.002 \text{ s}^{-1}$ | [55,56] |
| $R_{am} \rightarrow R_{ic} + \text{pheromone}$ | $k_{14}$ | $0.002 \text{ s}^{-1}$ | [42,43] |
| $R_{im} \rightarrow R_{ic}$ | $k_{15}$ | $0.0004 \text{ s}^{-1}$ | [42] |
| Diffusion coefficient of $R_{im}$ | $D_{\text{receptor}}$ | $0.0001 \mu\text{m}^2 \text{ s}^{-1}$ | [18] |
| Diffusion coefficient of pheromone | $D_{\text{pheromone}}$ | $150 \mu\text{m}^2 \text{s}^{-1}$ | [19] |

$R_{\text{im}}$ is membrane-bound inactive receptors. $\lambda_{11}$ is chosen to make sure the equilibrium dissociation constant ( $K_D$ ) for the receptor is 6-7 nM [56].

Although the actual pheromone gradients experienced by mating yeast cells have not been experimentally visualized, previous work simulated the pheromone gradient that would be perceived by a cell with an adjacent partner secreting pheromone from its polarity site [19] (Fig 5B), assuming experimentally measured pheromone emission rates [52]. To simulate cells that are initially searching for a partner, we initialized the simulations with a uniform pheromone concentration of 1.5 nM, which resulted in relocating Cdc42 clusters (S8 Fig). Then, to simulate a situation in which the cell orients polarity (and hence pheromone secretion) towards the partner cell, we imposed a pheromone gradient that varied between 1.5-5.8 nM (Fig 5C). In many but not all simulations, this gradient led to cessation of polarity site relocation within 50 min and development of a stable cluster of Cdc42 oriented towards the pheromone source (Fig 6A and S7 Movie). We next ran simulations using a gradient that ranged from 0-1.2 nM (Fig 5D). Remarkably, in this case all simulations produced a stable polarity site located in the region of high pheromone within 30 min (Fig 6B and S8 Movie). These results are consistent with experimental findings in which cells secreting only 20% of the amount of pheromone as wildtype cells are able to stabilize the polarity sites of mating partners [53]. Polarity stabilization occurred more rapidly in 0-1.2 nM than in 1.5-5.8 nM gradients (Fig 6C), consistent with the idea that the main obstacle to polarity stabilization is provided by competition with distinct polarity clusters seeded by the background levels of pheromone. In summary, our simulations of the combined polarity circuit demonstrate that while a uniform concentration of pheromone destabilizes polarity and promotes relocation, a local pheromone gradient can stabilize polarity.

**Fig 6.**
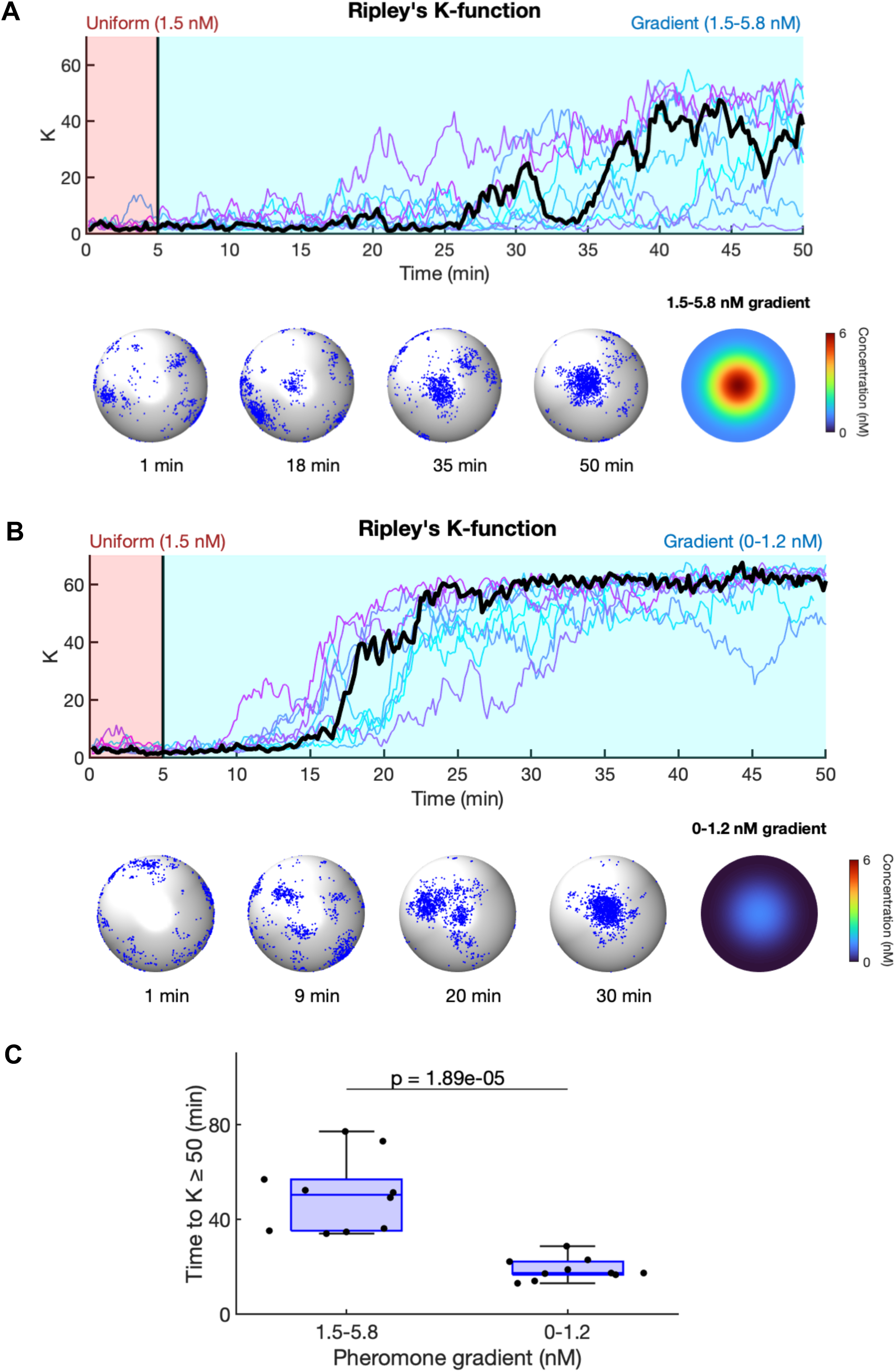
Pheromone gradients stabilize the polarity site. Simulation results using the two gradients shown in Fig 5. For both cases, cells were exposed to 1.5 nM uniform pheromone for the first 5 min. The pheromone gradient was applied starting at 5 min. **A**) Results for the gradient shown in Fig 5C. Time series for *K* for 10 simulations (Top). Distributions for active Cdc42 corresponding to the time series shown in black (Bottom). **B)** Same as **A** using gradient shown in Fig 5D. **C)** Time for *K* to 50 for the two gradients.

## Discussion

### Models of the yeast polarity circuit

The first realistic model for the yeast polarity circuit was proposed by Goryachev and Pokhilko, who considered a deterministic model that consisted of a set of reaction-diffusion equations [12]. They used their model to demonstrate that Bem1-GEF mediated positive feedback coupled with diffusion rates that vary between the membrane and cytosol was sufficient to generate a stable polarity site through a Turing mechanism [12]. Several more abstract minimalistic models, containing as few as two chemical species with non-linear positive feedback, were also studied to gain a deeper understanding of how cells might use Rho-family GTPases to generate spatial patterns. Because of their mathematical tractability, these models produced important insights into the mechanisms underlying establishment, competition and coexistence of GTPase clusters [14,15,30,57–61]. While theoretical work has begun to establish conditions that support polarity establishment and coarsening, less attention has been given to identifying mechanisms that produce movement or relocation of the polarity site.

Dynamic polarity site relocation is critical during the indecisive phase of yeast mating, when polarity clusters undergoes erratic assembly, disassembly, and seemingly random motion. Such behavior is unlikely to be captured by deterministic models, which ignore the molecular nature of biochemical systems and therefore miss noise-driven phenomena. In contrast, stochastic models attempt to capture stochastic effects that arise from the random nature of biochemical reactions and typically track molecule numbers as opposed to concentrations. Such particle-based simulations of a very simple two-species model were used to investigate the behavior of a system with linear positive feedback [30,31]. With linear positive feedback, clustering of Cdc42 only occurs in regimes with small molecule numbers, making the degree and location of clustering susceptible to molecular fluctuations. Hegemann et al. exploited the noise-driven relocation of polarity clusters in this model to explore how pheromone gradients might direct polarity to the correct site via the Far1 pathway [27]. However, experimental findings suggest that yeast polarization is robust to increased molecular abundances [16,62,63], indicative of more robust nonlinear positive feedback. Particle-based models with non-linear positive feedback could also display noise-driven relocation of polarity clusters, but (as with the linear feedback model) such behavior required fine-tuning and only occurred in a small region of parameter space [32]. Thus, it remained unclear whether simply incorporating molecular noise into a polarity circuit would suffice to explain the indecisive polarity behavior seen in mating yeast.

### Complementary positive feedback loops ensure successful mating

Successful mating requires that yeast cells direct their growth toward potential mating partners, which requires proper positioning of the polarity site. To achieve this task, the polarity system must satisfy several often competing requirements: 1) detection of pheromone gradients, 2) establishment of a single polarity site, 3) reorientation of the polarity site when misaligned with the gradient, and 4) stabilization of the polarity site when oriented toward a mating partner. Our modeling results suggest that these requirements are met through a series of three coupled positive feedback loops acting on different time scales (Fig 7).

**Fig 7.**
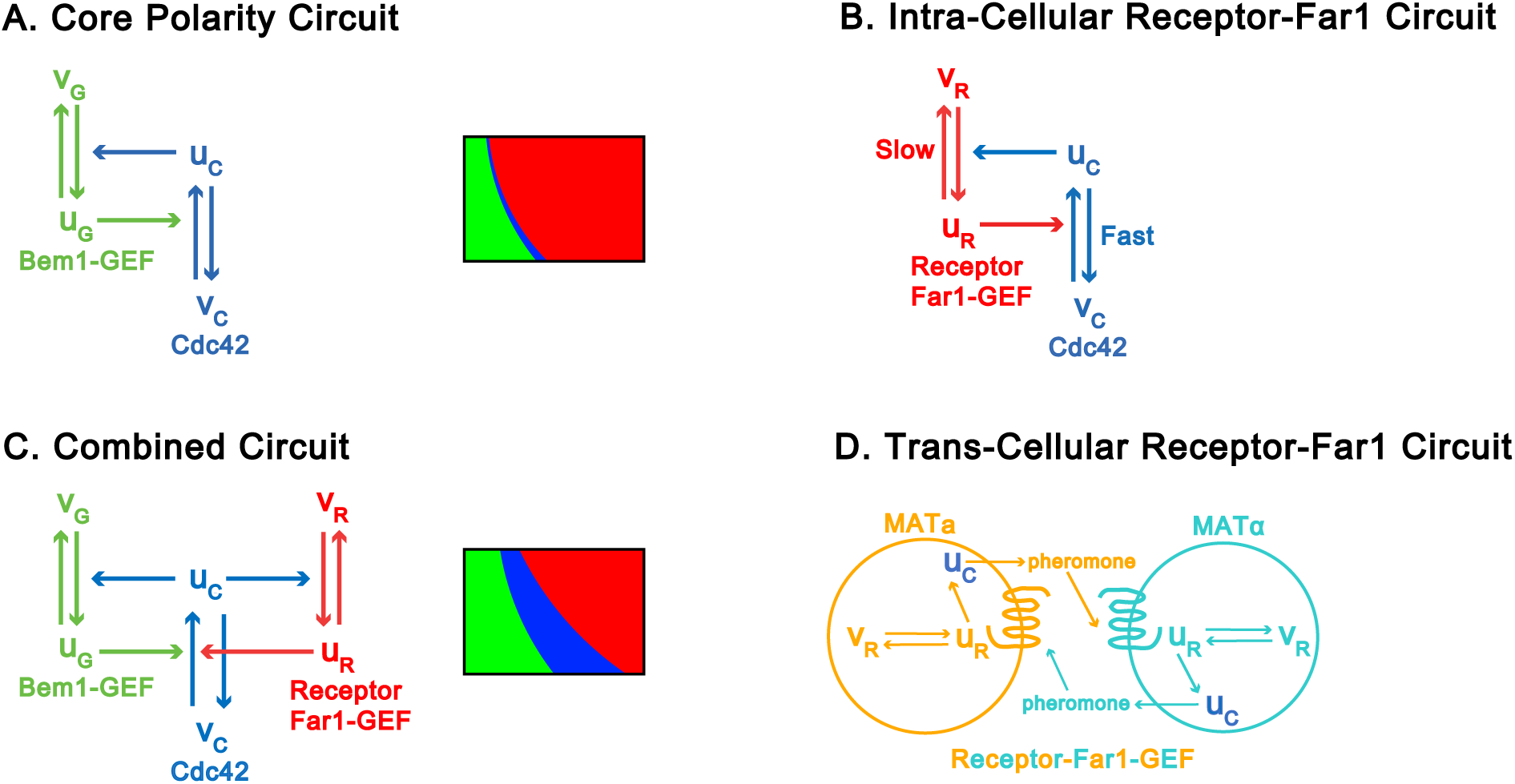
Summary of system architecture. Schematics of positive feedback loops for **A)** the core polarity circuit, **B)** the intracellular receptor-Far1 circuit, **C)** the combined circuit and **D)** the trans-cellular receptor-Far1 circuit. *u* denotes the membrane-associated slowly-diffusing “active” form, while *v* denotes the cytosolic, rapidly diffusing “inactive” form of the relevant protein—Cdc42 is indicated by the subscript C, Bem1-GEF by the subscript G, and Receptor-Far1-GEF by the subscript R.

The first positive feedback loop forms the core polarity circuit (Fig 7A). In this circuit, positive feedback occurs because active Cdc42 molecules increase the activation rate of neighboring Cdc42 molecules by recruiting active GEF molecules to their location. Because diffusion in the cytosol is fast as compared to diffusion in the membrane, this positive feedback loop provides an effective mechanism for forming clusters of active Cdc42. Rapid diffusion in the cytosol also allows competition between coexisting clusters for polarity factors in the cytosol. Larger clusters generate stronger positive feedback and thus are able to out-compete smaller clusters, leading to a “winner take all” mechanism for establishing a single polarity site.

While strong positive feedback is effective at generating a unique polarity site, it also exhibits undesirable features for gradient tracking. First, initiation of a polarity cluster is highly susceptible to fluctuations in the local concentrations of relevant molecules, so that polarity sites can easily form at sites that are misaligned with the pheromone gradient. Second, it is difficult to relocate a polarity site once it has been established: except in a very narrow slice of parameter space, this makes polarity sites resistant to molecular fluctuations (Fig 7A). The Far1 pathway by which the polarity circuit is coupled to pheromone sensing in yeast appears to overcome both of these drawbacks (Fig 7B). Active pheromone receptors recruit GEF molecules in complex with the multifunctional protein Far1, thereby activating Cdc42 near active receptors. Thus, active receptors continuously seed formation of new Cdc42 clusters, while blunting competition between coexisting clusters. This diminished competition generates dynamic clusters of Cdc42 that form and disassemble throughout the cell surface, providing a search mechanism for finding a local pheromone source. The combined polarity/Far1 circuit exhibits a much larger region of parameter space where molecular fluctuations can drive polarity site relocation (Fig 7C).

Interestingly, receptor-mediated activation of Cdc42 forms a second positive feedback loop because active Cdc42 molecules recruit actin cables along which vesicles containing new receptors that are trafficked (Fig 7B). Due to long receptor lifetimes on the membrane [64,65], this feedback loop does not support polarity establishment under conditions of constant pheromone concentration. However, in the presence of a pheromone gradient, positive feedback by the receptor-Far1 circuit operates more strongly in the region of high pheromone concentration, because receptor activation is higher leading to more active Cdc42 in this region. In turn, higher Cdc42 activity leads to an increase in the local concentration of receptors, reinforcing Cdc42 activity and promoting the formation of a stable polarity site. Interestingly, this mechanism of gradient tracking is able to respond effectively to pheromone gradients of low peak concentration (1.2 nM), consistent with observations that cells in which pheromone production has been reduced by 80% are still able to successfully signal their location to mating partners (Jacobs 2022). A potential limitation of this mechanism for gradient tracking is that cells can be confused when the gradient is superimposed on a background of constant pheromone (e.g., the case in which the gradient ranges from 1.5-5.8 nM). As pheromone concentrations experienced by yeast in the wild are likely to be quite variable, we speculate that yeast have evolved additional mechanisms to overcome this limitation.

Finally, a third positive feedback loop acts between cells of opposite mating type (Fig 7D). All haploid yeast cells emit mating-type-specific pheromones, which bind to receptors on the surface of cells of the opposite mating type. Signaling through these receptors both increases the rate of pheromone production within the receiving cell and, by locally activating Cdc42, increases the probability that pheromone is released in the direction of the opposite mating type. Our simulations confirm experimental findings that this localized release of pheromone is important for establishing gradients steep enough to be detected by the adjacent mating partner [19]. This positive feedback loop is likely to play a role during the indecisive phase of mating when both cells’ polarity sites are dynamic. Considering this two-cell scenario will be the subject of future investigations.

### Biological implications and future directions

Our conclusions may be applicable to polarity regulation in other systems. Similar to *S. cerevisiae*, polarity sites in the distantly related fission yeast *S. pombe* also exhibit erratic assembly-disassembly behavior during mating, and are stabilized when they happen to align with a partner [66,67]. Interestingly, fission yeast lack the Far1 pathway, but possess a pathway with the same architecture, where pheromone-receptor binding leads to local recruitment of a Cdc42 GEF [68]. Thus, our findings may apply broadly to fungal mating systems [25].

Although our work suggests that molecular noise is sufficient to promote relocation of polarity clusters in cells that combine the polarity circuit with the receptor-Far1 pathway, other phenomena may also promote mobility of polarity clusters. In particular, polarized vesicle trafficking can cause dilution and lateral displacement of polarity factors [55,69–71]. In the future, it may be useful to investigate whether a model combining vesicle traffic and the receptor-Far1 pathway can accurately reproduce polarity site behaviors in mating yeast.

Our proposed mechanism of polarity site relocation has implications for a range of cell functions beyond mating yeast cells. For example, migratory cells often exhibit relocating polarity sites while moving in chemical gradients [1,72,73]. In axon development, multiple polarity sites (neurites) are formed initially, and one of them differentiates into an axon as it moves up chemical gradients [74]. Many of these functions require receptor endocytosis [75–77]. It will be interesting to investigate whether dynamic polarity might result from differences in the time scales of polarity factor diffusion and receptor trafficking. Comparing the principles of polarity in mating yeast to those in other eukaryotic cells promises to provide insight into many cell functions.

## Methods

### 2D Particle-based model

All particle-based simulations were performed using Smoldyn (v2.67) with continuous space and discretized time intervals on a Linux-based computing system (Longleaf cluster at UNC Chapel Hill, 2.50 GHz and 2.30 GHz Intel Processors) [35,36]. Periodic boundary conditions were assumed in both spatial directions. Molecules were regarded as point particles with no volumes. Brownian motion of the molecules was simulated with the Euler-Maruyama method as follows. Let *x*(*t*) and *y*(*t*) represent the coordinates of a given molecule at time *t*, then molecule’s position at *t +* Δ*t* is calculated as:

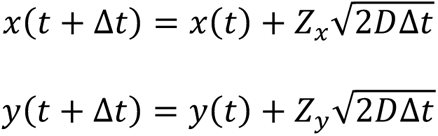

where *Z_x_* and *Z_y_* are independent random numbers drawn from the standard normal distribution, and *D* is the diffusion coefficient. Coordinates of all species were stored every 10 seconds from a 4000-second simulation.

Second-order reactions are governed by two parameters: the reaction radius ρ and reaction rate λ. When two reacting partners are within a distance ρ, then the probability that they react within a time step Δ*t* is *P* = 1-exp(-λΔ*t*). When Δ*t* is sufficiently small, *P* ≈ λΔ*t.* When bound molecules dissociate, they are placed at a distance which is 0.00001 μm beyond the binding radius ρ to avoid immediate reassociation. First-order reactions occur with the probability *P* = 1-exp(-*k*Δ*t*), where *k* is the rate constant for the reaction. Parameter values used in the simulations are listed in Tables 1-2.

### 3D Particle-based model

The yeast cell was approximated as a sphere. Membrane-bound species diffuse on the surface of the sphere. Cytosolic species diffuse inside the sphere. Reaction rates for the 3D model are listed in Tables 1-3. Initial choices for the rate constants in the 3D simulations were obtained by converting rate constants from the 2D simulations using the method of Ramirez et al. [32]. However, a few rate constants required additional tuning to ensure the steady-state concentrations of the 2D and 3D simulations were similar. Rate constants for reactions between membrane-bound species did not change between the 2D and 3D simulations. Rate constants for first-order reactions in which a cytosolic species associated with the membrane were scaled as follows:

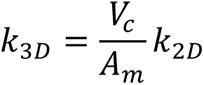

*V_c_* is the volume of the sphere, and *A_m_* is the area of the membrane.

The rate constants for second-order reactions that involved a cytosolic species and membrane-bound species were scaled as follows:

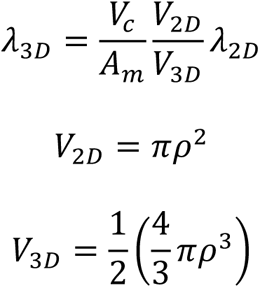

*V_2D_* is the 2D reaction area, and *V_3D_* is the reaction volume for 3D simulations. The reason *V_3D_* is half the volume of a sphere is because one of the reactants is membrane-bound.

### Simulating receptor cycling

Actin cables directed by Cdc42 bring pheromone receptors to the polarity site. We did not explicitly take into account actin cables in our simulations. Actin-dependent delivery of receptors was implicitly modeled as a bimolecular reaction. Each individual Cdc42-GTP is able to recruit a receptor from the cytoplasm to the membrane with a specified rate (λ_9_ in Table 2, and λ_12_ in Table 3) during each time step. Membrane receptors are internalized via endocytosis, which was modeled as a first-order reaction with specified rate constants (k_10_ in Table 2, and k_14_, k_15_ in Table 3).

### Simulating pheromone molecules

3D Pheromone gradients were simulated according to the method described previously [19]. A pair of mating cells were modeled as two spheres with a diameter of 5 μm. One cell emitted pheromone. We assumed that the number of released pheromone molecules followed a Poisson process, and all molecules were emitted from a single point source. Vesicle release events were simulated in Smoldyn using the command *pointsource*. Pheromone was removed at a spherical absorbing boundary 7 μm from the origin. Cell membranes were treated as reflecting boundaries.

### Quantifying polarity

Polarity was measured using normalized versions of Ripley’s K-function [17,32,40,41,78]. The function *K*(*r*) measures the deviation of the current particle distribution from a uniform distribution based on the cumulative distribution of pairwise molecular distances *P*(*r*).:

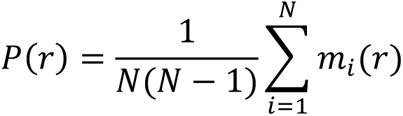

For particle distributions on a 2D plane:

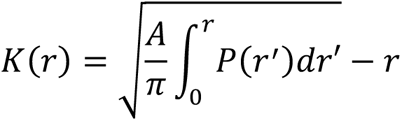

For particle distributions on a 3D sphere:

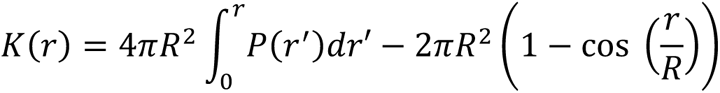

where *m_i_*(*r*)Δ*r* is the number of molecules at a distance *d* from molecule *i* such that *r* ≤ *d ≤ r+*Δ*r*, *N* is the total number of molecules in the domain, and *A* is the area of the domain. *K*(*r*) = 0 if particles are distributed uniformly. For a non-uniform distribution, the K-function typically goes through a maximum value for a finite value of *r*. To quantify polarity, we used the maximum value of *K*. For 2D simulations, we defined a molecule distribution with *K* < 1.5 as unpolarized states; *K* ≥ 1.5 as polarized states; and *K* ≥ 3 as stable polarity (S1A Fig). For 3D simulations, we defined a distribution with *K* < 30 as unpolarized states; *K* ≥ 30 as polarized states; and *K* ≥ 50 as stable polarity (S1B Fig).

### Identifying polarity regimes

To identify the three polarity regimes (unpolarized, transient polarity, and polarized), we recorded values of *K* every 10 seconds during the last 2000 seconds of each simulation. Simulations were run using two kinds of initial conditions: random and polarized. For random initial conditions, positions of the Cdc42 molecules were chosen from a uniform distribution, whereas for the polarized case position for the Cdc42 molecules were chosen according to a Gaussian in the center of the domain with a standard deviation of 0.2. For a given number of Cdc42 and Bem1-GEF molecules, we ran ten simulations for each type of initial condition. For each simulation, we calculated the difference Δ between the number of polarized states *F*(*K* ≥ 1.5) and number of unpolarized states *F*(*K* < 1.5):

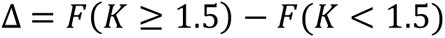

If the system spends half its time in a polarized state and half unpolarized, Δ equals 0. A large positive value of Δ indicates a polarized regime and a large negative value indicates an unpolarized regime. Let *N* equal the total number of *K* values recorded during simulation. To identify the transient polarity regime, we used the condition:

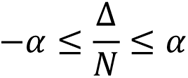

with α = 0.85. If Δ*/N* > α, the regime was categorized as polarized and Δ*/N* < -α unpolarized. For a given number of Cdc42 molecules, the relationship between Δ and the number of Bem1-GEF molecules, *x*, was assumed to follow the logistic function:

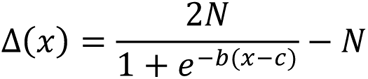

This function was then fit to the simulation results to determine values of Bem1-GEF molecules at which the system transitioned between polarity regimes (S2 and S4 Figs).

### Cluster detection by Voronoi tessellation

We utilized a previous method based on Voronoi tessellation to detect clusters [48,79]. A cluster was defined as a group of adjacent Voronoi polygons with sizes below a predetermined threshold. The threshold for polygon sizes was determined as follows. As a negative control, we generated 50 realizations of particle distributions from a uniform distribution. These distributions were used to generate a probability density for Voronoi polygon sizes. We also obtained the probability density for the polygon size distribution of Cdc42-GTP molecules generated from our simulations. The intersection of the probability densities from the negative control and our simulations defines the threshold of polygon sizes for cluster formation. A polygon was deleted from the Voronoi diagram if its size was above the threshold, meaning that the density of molecules in the polygon was too low to be considered as part of a cluster. The remaining polygons on the Voronoi diagram would be considered as components of potential clusters. Then, we grouped the neighboring polygons into clusters and applied filters for cluster area and number density. The threshold area of a cluster was set to be 0.785 μm^2^ as the diameter of a polarity site of yeast was estimated to be 1 μm. We set the threshold number of Cdc42 in a cluster according to a previous measurement [62]. Groups of polygons passed the filters were recorded as true clusters.

## Data availability

The code used to generate figures and simulations is publicly available at https://github.com/guanhighyun/Indecisive-polarity-behavior

## Acknowledgements

We thank Steven S. Andrews for developing and updating the computer program Smoldyn which is used for simulations in the project. We also thank all members of the Elston lab for helpful discussions on the project.

## Supporting information

**S1 Fig. Examples of the Ripley’s K-function**. Cluster distributions corresponding to the listed *K* values. **A)** 2D examples for a square domain of length 8.8623 μm. **B)** 3D examples for a sphere of radius 2.5 μm.

**S2 Fig. Polarity regimes for the core polarity circuit. A)** Difference in the number of polarized and unpolarized states as a function of Bem1-GEF abundance. Data points (o) represent simulation results for the total number of Cdc42 molecules indicated in the plot title. Curves represent fits to the data using a logistic function (unpolarized regime - green, polarized regime - red, and transient regime - blue). **B** and **C)** Distributions for *K* in the transient regime using a total Cdc42 abundance of 3000. Stars indicate the Bem1-GEF abundances used to produce the distributions (100 lower panels and 106 upper panels).

**S3 Fig. Simulation results for unpolarized and polarized regimes of the core polarity circuit**. **A)** Time series for K for 10 simulations in the unpolarized regime (left panel). Distributions for active Cdc42 taken form black time series using time points indicated with ticks on the time axis (right panels). Simulations were performed using uniform random distributions of molecules as initial conditions with 3000 Cdc42 and 70 Bem1-GEF molecules. **B)** Same as A except with 3000 Cdc42 and 120 Bem1-GEF molecules.

**S4 Fig. Polarity regimes for the combined polarity circuit. A)** Difference in the number of polarized and unpolarized states as a function of Bem1-GEF abundance. Data points (o) represent simulation results for the total number of Cdc42 molecules indicated in the plot title. Curves represent fits to the data using a logistic function (unpolarized regime - green, polarized regime - red, and transient regime - blue). **B** and **C)** Distributions for *K* in the transient regime using a total Cdc42 abundance of 3500. Stars indicate the Bem1-GEF abundances (135 lower panels and 165 upper panels). All simulations were performed using 30 Far1-GEF and 2500 receptor molecules.

**S5 Fig. Simulation results for unpolarized and polarized regimes of the combined polarity circuit. A)** Time series for K for 10 simulations in the unpolarized regime (left panel). Distributions for active Cdc42 taken form black time series using time points indicated with ticks on the time axis (right panels). Simulations were performed using uniform random distributions of molecules as initial conditions with 3000 Cdc42, 80 Bem1-GEF, 30 Far1-GEF and 2500 receptor molecules. **B)** Same as A except with 280 Bem1-GEF.

**S6 Fig. Number of clusters as a function of time. A)** Time series for the number of clusters for 10 simulations of the core polarity circuit with 3000 Cdc42 and 102 Bem1-GEF molecules. Results shown in black highlight a single time series. **B)** Same as A except using the combined polarity circuit model with 3000 Cdc42, 170 Bem1-GEF, 30 Far1-GEF molecules and 2500 receptor molecules.

**S7 Fig. Comparison of 2D and 3D models.** Simulations were performed with 3000 Cdc42, 170 Bem1-GEF, 30 Far1-GEF, and 2500 receptor molecules. Simulations started with uniformly distributed molecules. Lines represent molecule number averages and the shaded areas represent ± std for 30 simulations (3D – blue, 2D – red). All simulations were run for 66 min.

**S8 Fig. Indecisive polarity behavior with uniform pheromone concentration.** Simulations were performed using 3000 Cdc42, 170 Bem1-GEF, 2500 receptor, 30 Far1-GEF and 1.5 nM uniform concentration of pheromone molecules.

**S1 Movie. Spatiotemporal distribution of Cdc42-GTP in a transient polarity regime of the core polarity circuit model simulation**. Blue dots represent individual molecules of active Cdc42. Corresponds to Fig 1D.

**S2 Movie. Spatiotemporal distribution of Cdc42-GTP in a transient polarity regime of the combined polarity circuit model simulation.** Corresponds to Fig 3C.

**S3 Movie. A cluster of Cdc42-GTP dissipates after introducing the Far1 pathway**. Receptors and Far1-GEF molecules were introduced at 400 seconds. Corresponds to Fig 3D.

**S4 Movie. A cluster of Cdc42-GTP is established and maintained if receptors are initially polarized.** Corresponds to Fig 3E.

**S5 Movie. Cdc42-GTP clusters relocate between fixed Far1-GEF molecules.** 15 fixed Far1-GEF molecules directly activate Cdc42 without receptors. Corresponds to Fig 4E.

**S6 Movie. Spatiotemporal distribution of Cdc42-GTP of the 3D combined polarity circuit model simulation.** Corresponds to Fig 4F.

**S7 Movie. A 1.5-5.8 nM pheromone gradient can promote stable formation of Cdc42-GTP clusters.** 1.5 nM uniform pheromone was applied for the first 5 min. Starting from 5 min, a gradient was imposed. Corresponds to Fig 6A.

**S8 Movie. A 0-1.2 nM pheromone gradient can efficiently stabilize Cdc42-GTP clusters.** 1.5 nM uniform pheromone was applied for the first 5 min. Starting from 5 min, a gradient was imposed. Corresponds to Fig 6B.

