## Supplementary figures and images for "Particle-based simulations reveal two positive feedback loops allow relocation and stabilization of the polarity site during yeast mating"

### S1 Fig

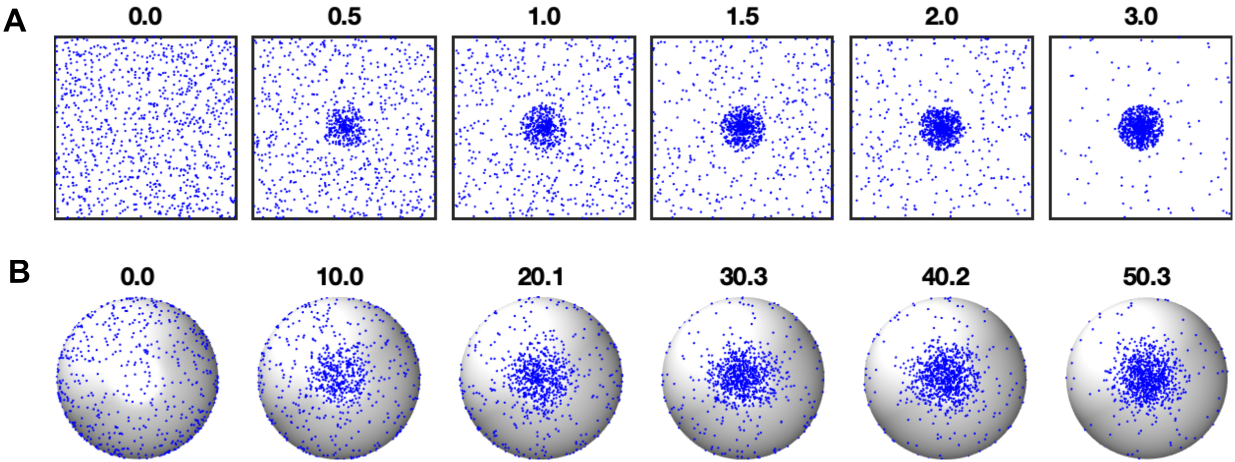

### S2 Fig

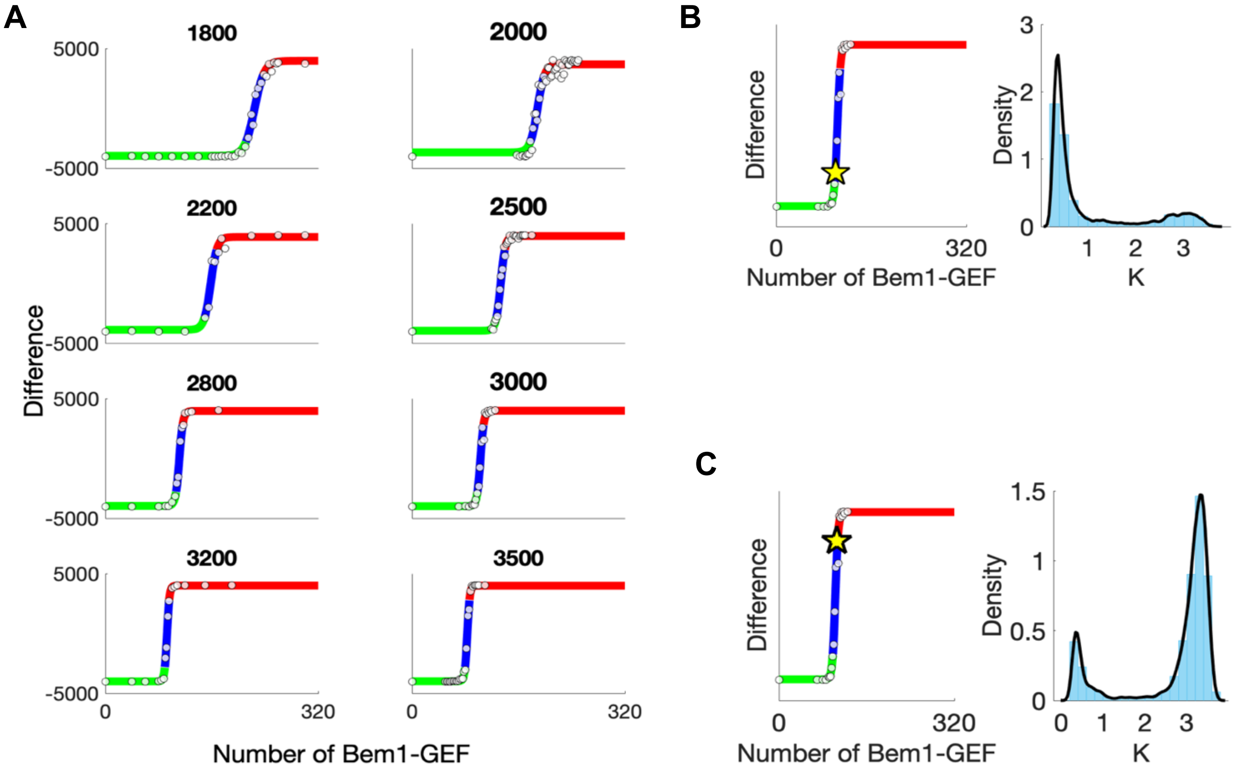

### S3 Fig

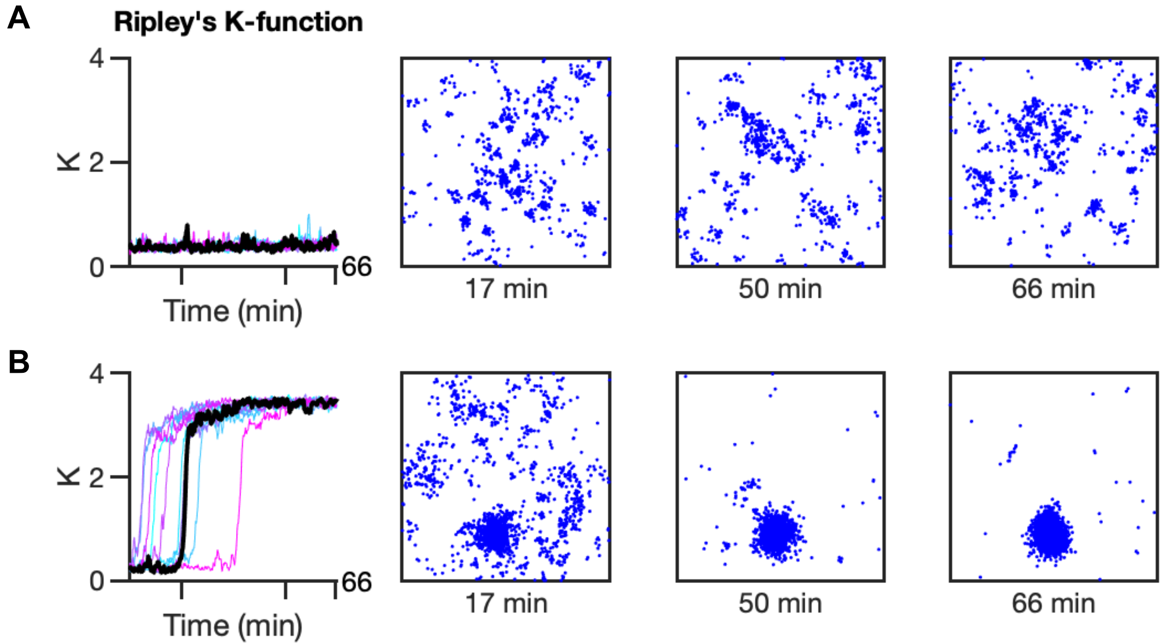

### S4 Fig

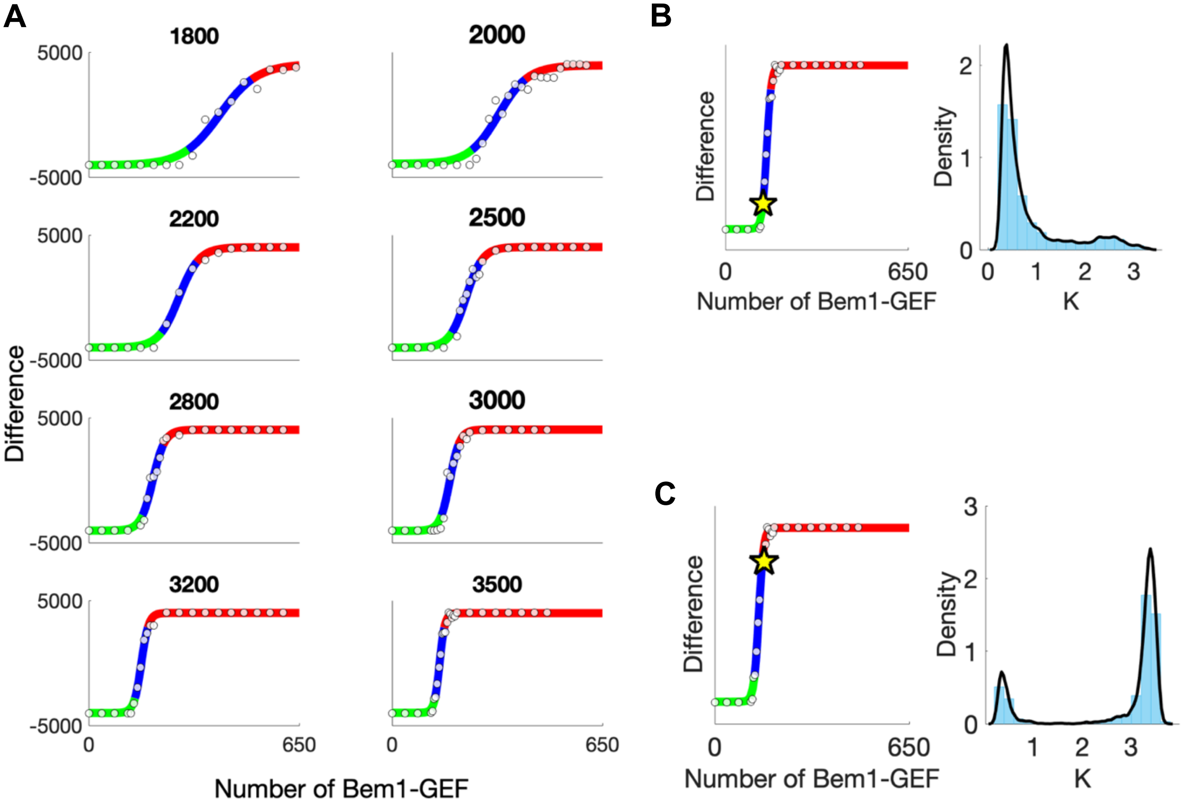

### S5 Fig

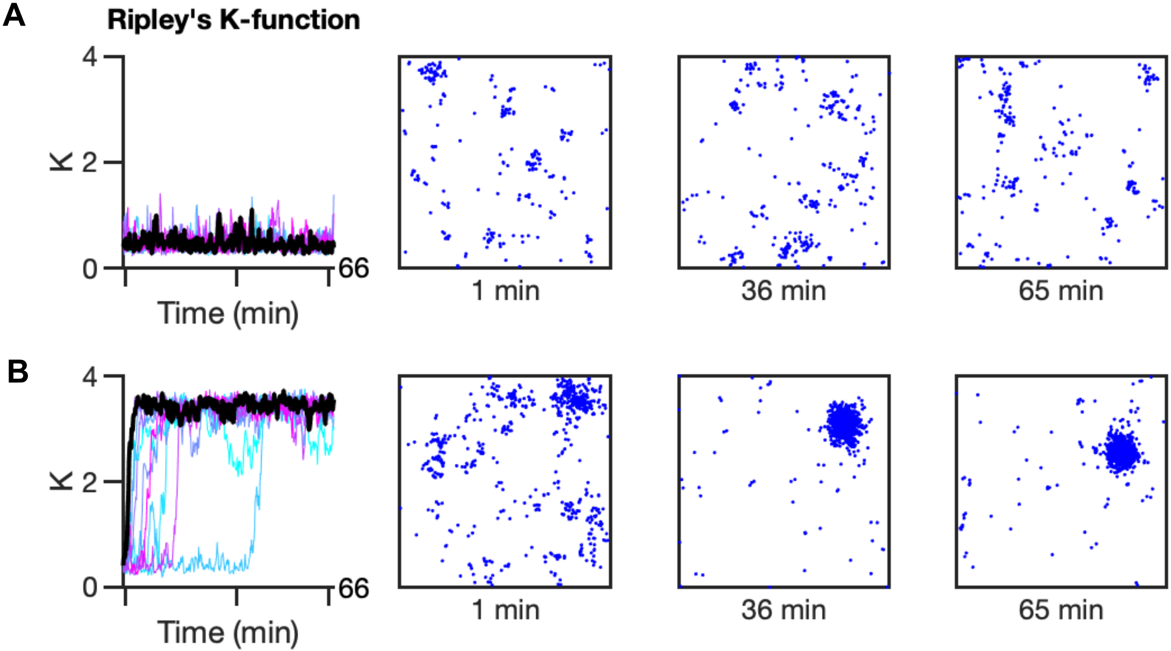

### S6 Fig

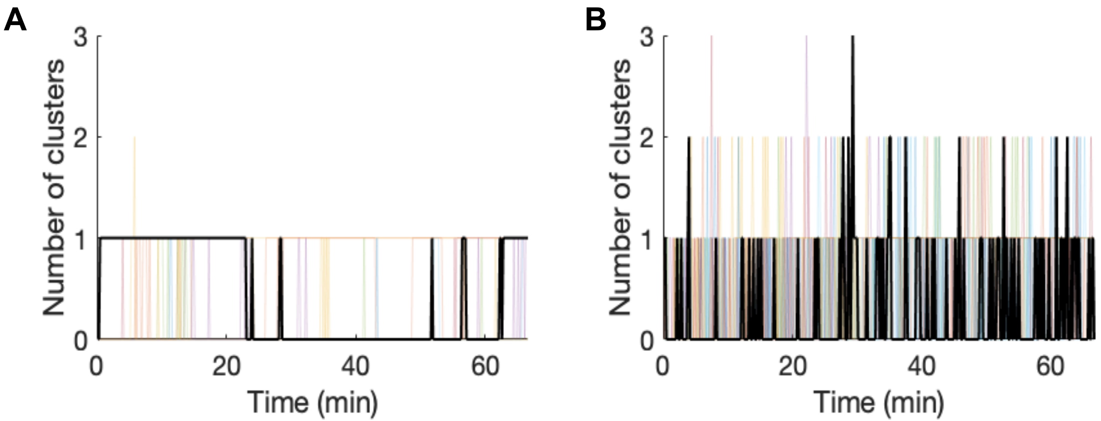

### S7 Fig

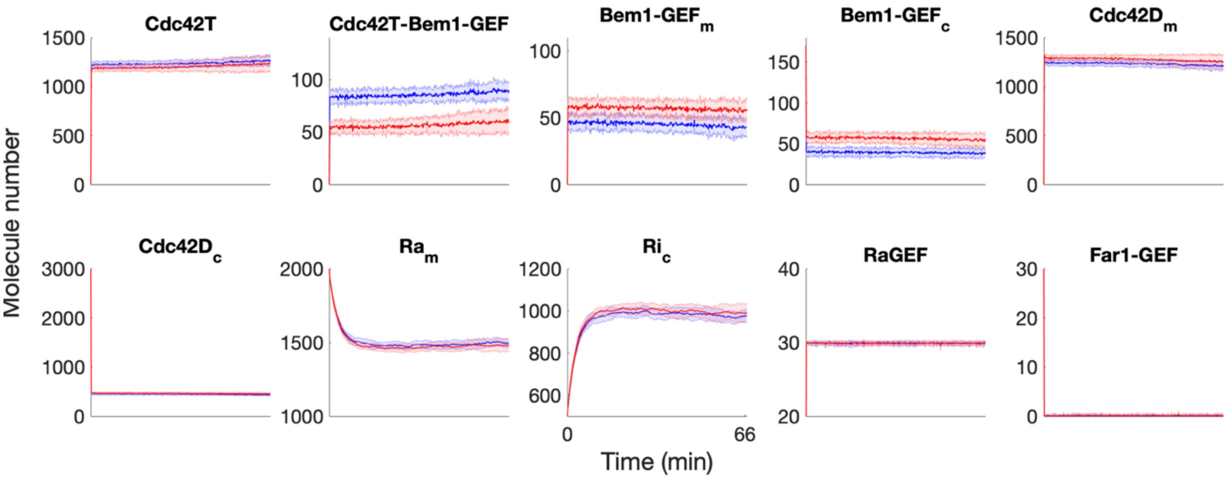

### S8 Fig

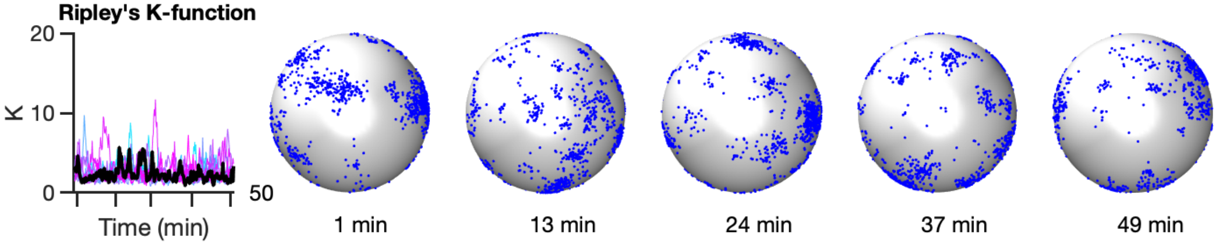
